# Genetic dissection of regulation by a repressing and novel activating corrinoid riboswitch enables engineering of synthetic riboswitches

**DOI:** 10.1101/2023.06.26.546531

**Authors:** Rebecca R. Procknow, Kristopher J. Kennedy, Maxwell Kluba, Lesley J. Rodriguez, Michiko E. Taga

**Affiliations:** Department of Plant & Microbial Biology, University of California Berkeley, Berkeley, CA USA

## Abstract

The ability to sense and respond to intracellular metabolite levels enables cells to adapt to environmental conditions. Many prokaryotes use riboswitches – structured RNA elements usually located in the 5’ untranslated region of mRNAs – to sense intracellular metabolites and respond by modulating gene expression. The corrinoid riboswitch class, which responds to adenosylcobalamin (coenzyme B_12_) and related metabolites, is among the most widespread in bacteria. The structural elements for corrinoid binding and the requirement for a kissing loop interaction between the aptamer and expression platform domains have been established for several corrinoid riboswitches. However, the conformational changes in the expression platform that modulate gene expression in response to corrinoid binding remain unknown. Here, we employ an *in vivo* GFP reporter system in *Bacillus subtilis* to define alternative secondary structures in the expression platform of a corrinoid riboswitch from *Priestia megaterium* by disrupting and restoring base-pairing interactions. Moreover, we report the discovery and characterization of the first riboswitch known to activate gene expression in response to corrinoids. In both cases, mutually exclusive RNA secondary structures are responsible for promoting or preventing the formation of an intrinsic transcription terminator in response to the corrinoid binding state of the aptamer domain. Knowledge of these regulatory mechanisms allowed us to develop synthetic corrinoid riboswitches that convert repressing riboswitches to riboswitches that robustly induce gene expression in response to corrinoids. Due to their high expression levels, low background, and over 100-fold level of induction, these synthetic riboswitches have potential use as biosensors or genetic tools.

## Introduction

Organisms rely on gene regulation to direct resources toward the physiological needs of the moment. Metabolites are often sensed via metabolite-binding receptor proteins, but bacteria also sense and respond to metabolites using metabolite-binding RNAs known as riboswitches (1, 2). Riboswitches are structured RNAs, usually located in the 5’ untranslated region (UTR) of mRNAs, that change conformation to either promote or prevent gene expression in response to direct binding of an effector (3). They often regulate genes related to the synthesis, transport, or use of the effector to which they respond (4). Since their discovery in 2002, over 50 riboswitch classes have been characterized that respond to a range of effectors including amino acids, metal ions, nucleotides, and vitamins (3, 5–10). In addition to their natural forms, synthetic riboswitches have also been developed for use as biological tools (11, 12).

All riboswitches have two domains that communicate with each other via the formation of alternative secondary structures. Binding of the effector to the aptamer domain induces the formation of secondary structures in the expression platform domain that influence either translation or transcription elongation of downstream genes (2). In translational riboswitches, a hairpin can form in the expression platform that sequesters the ribosome binding site (RBS) to prevent translation, while the expression platform in transcriptional riboswitches can form an intrinsic transcription terminator hairpin. Most known riboswitches downregulate gene expression in response to effector binding (repressing riboswitches), but some have been found to induce expression (activating riboswitches).

The corrinoid riboswitch class (originally named adenosylcobalamin, cobalamin, or B_12_-riboswitches) is among the most widespread in prokaryotic genomes (3, 13). All known corrinoid riboswitches repress translation or transcription of genes for cobalamin biosynthesis, uptake, cobalamin-independent isozymes, or other functions in response to cobalamin binding (14). Like other riboswitch classes, corrinoid riboswitches can distinguish between structurally similar metabolites such as different cobalamin forms containing either a large (deoxyadenosyl) or small (methyl or hydroxyl) upper axial ligand (15). However, unlike other riboswitch classes, the effectors for corrinoid riboswitches are a group of naturally occurring variants of cobalamin – corrinoids with variation in the lower axial ligand – and we previously found that corrinoid riboswitches can respond to multiple corrinoids (16).

Another unusual feature of corrinoid riboswitches is that they rely on a tertiary base-pairing interaction (kissing loop) between loops L5 of the aptamer domain and L13 of the expression platform for effector sensing and repression of gene expression upon cobalamin binding (17, 18). Previous studies of the *E. coli btuB* and *env8*HyCbl riboswitches demonstrated that the kissing loop interaction modulates the formation of the RBS hairpin to prevent translation (17, 18). Specifically, kissing loop formation stabilizes the P13 stem, which promotes the formation of the RBS hairpin, while translation occurs when P13 formation is not stabilized by the kissing loop (18). X-ray crystal structures of translational and transcriptional corrinoid riboswitches resolve the effector-bound states, often including a kissing loop, but these crystal structures do not include other parts of the expression platform such as the RBS hairpin or terminator (15, 19, 20). It is not known how the effector-binding state promotes the formation of alternative secondary structures in the expression platform, leading to inhibition of translation or transcription (21). It is also unknown whether the kissing loop modulates the formation of the terminator hairpin in transcriptional riboswitches, as most prior studies focused on corrinoid binding and structural conformations in translational riboswitches (20).

Here, we have determined how the effector binding state of the aptamer domain of a model corrinoid riboswitch triggers the formation of alternative RNA structures in the expression platform. Whereas previous studies primarily relied on *in vitro* biochemical and structural approaches, the present study examines the regulatory mechanisms of transcriptional corrinoid riboswitches using an *in vivo* approach, which enabled us to measure the impacts of dozens of mutant riboswitches on regulation in an intracellular context. By constructing targeted mutations predicted to disrupt and restore base-pairing interactions in the expression platform of the *Priestia* (formerly *Bacillus*) *megaterium metE* riboswitch, we identified two alternative structural states in the expression platform that couple corrinoid detection to transcription. We additionally present the discovery of the first known corrinoid riboswitch that activates gene expression in response to corrinoid binding and identify the alternative structural states involved in its corrinoid response. Studying a repressing and an activating riboswitch allowed us to apply the ‘rules’ of the two regulatory strategies to flip the regulatory sign of the repressing riboswitch to create synthetic riboswitches that activate gene expression in response to cobalamin. Some of these synthetic activating riboswitches have a higher maximum expression and fold change than the natural activating riboswitch and could be used as corrinoid-detecting biosensors or regulatory systems.

## Results

### Model for corrinoid-responsive regulation in the *P. megaterium metE* riboswitch

We chose to dissect the regulatory mechanism of the *P. megaterium metE* riboswitch due to its high fold repression (26-fold) in our *B. subtilis* GFP reporter system (16). This riboswitch downregulates GFP expression in response to cobalamin binding in a dose-dependent manner (Figure 1A) and is predicted to be a transcriptional riboswitch (16). We developed a model for the formation of competing structures in the expression platform based on predicted secondary structures in the expression platform (Figure 1B and Figure S1A and B) (13, 22). According to this model, a kissing loop forms between L5 and L13 when the aptamer domain is bound to a corrinoid. The P13 stem, when stabilized by the kissing loop, is predicted to promote the formation of a terminator hairpin. In the absence of corrinoid binding, we predict that a portion of the 3’ side of the P13 stem pairs with part of the 5’ side of the terminator stem, forming an antiterminator that prevents the formation of the terminator hairpin. This model contrasts with models of other corrinoid riboswitch expression platforms, which are proposed to form alternative structures with bases from the aptamer domain or other portions of the expression platform (18–20). To test different aspects of the model, we disrupted and restored Watson-Crick-Franklin complementary base-pairing interactions predicted to stabilize one of the two predicted conformations.

**Figure 1.**
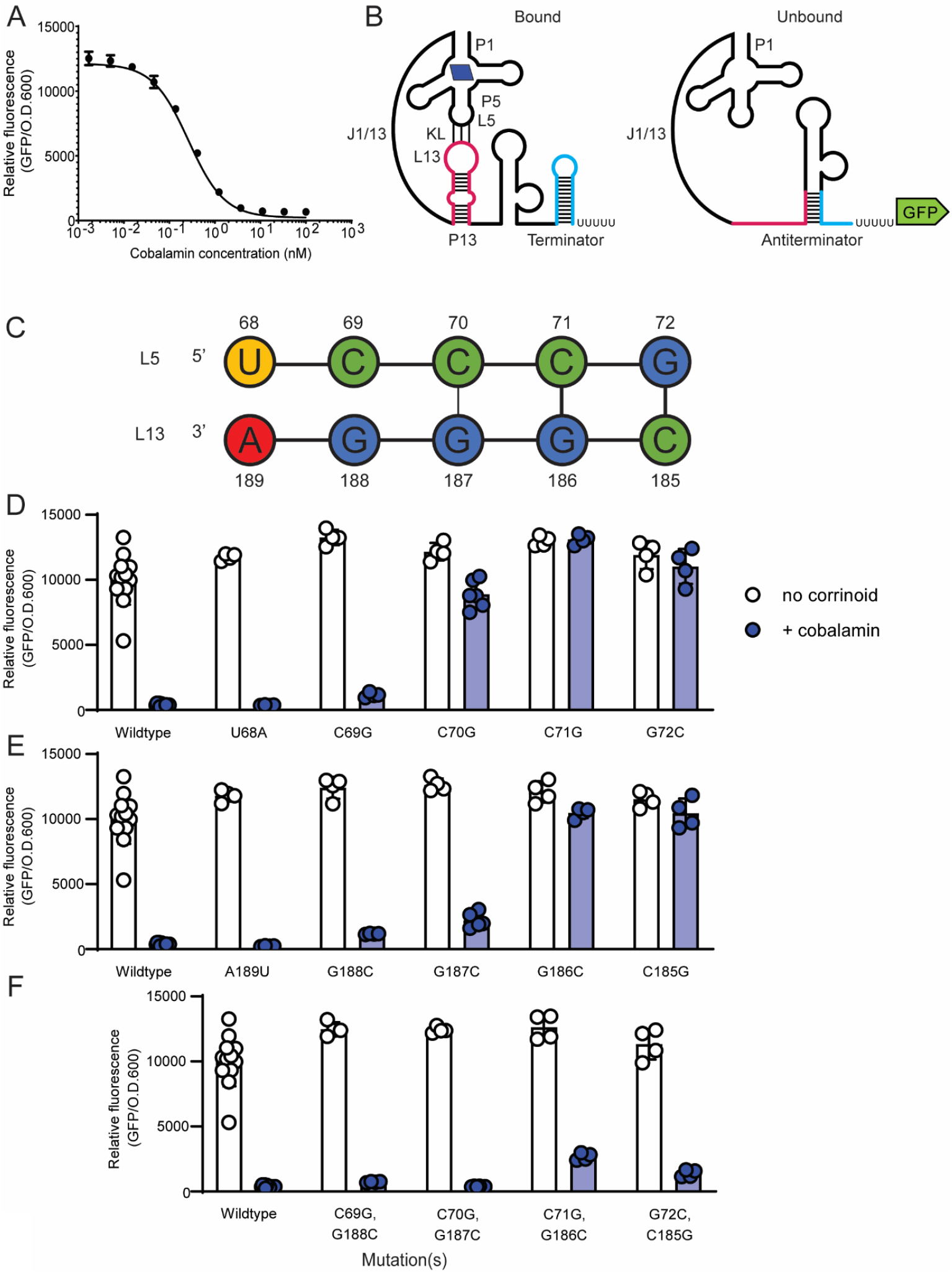
Model for the regulatory mechanism of the repressing *P. megaterium metE* riboswitch and dissection of the kissing loop. (A) Dose response of the *P. megaterium metE* riboswitch to cobalamin in the *B. subtilis* GFP reporter. (B) A model for the effector-bound (left) and unbound (right) conformations of the riboswitch. The effector-bound state is depicted with cobalamin (blue parallelogram) and the kissing loop (KL) interaction between loop (L) 5 and L13. Bases belonging to the paired stem (P) 13 and terminator hairpins are depicted as pink or blue in both structures, respectively. (C) The wildtype kissing loop sequence. Base numbers are relative to the first base in the P1 stem. The influence of point mutations in (D) L5, (E) L13, or (F) combined point mutations in L5 and L13 meant to restore the kissing loop interaction was measured in the *B. subtilis* GFP reporter system without (white) or with (blue) addition of 100 nM cobalamin. Genotypes are listed in Table S1. Data from four or more biological replicates are shown; bars and error bars represent mean and standard deviation, respectively.

### Mutational analysis defines the kissing loop in the *P. megaterium metE* riboswitch

To determine the mechanism of regulation in the *P. megaterium metE* riboswitch, we first mutated each base in the predicted kissing loop and measured its impact on GFP expression in the *B. subtilis* reporter assay. The sequence of L5 exactly matches the UCCCG consensus defined by McCown et al. 2017 (13), and the predicted L5-L13 kissing loop contains five contiguous and complementary base pairs (Figure 1C), as in the *E. coli btuB* translational riboswitch (18), but distinct from the *env8* translational riboswitch, which contains both a mismatch and a bulge (17).

We found that some of the mutations in L5 and L13 of the *P. megaterium metE* riboswitch result in constitutive GFP expression, indicating a disruption in the ability to sense and respond to corrinoid, while other mutations have little or no impact on expression despite all L5 bases being highly conserved (Figure 1D, E). According to these results, the base pairs in L5·L13 that are most involved in the kissing loop interaction are G72·C185, C71·G186 and, to a lesser extent, C70·G187. Mutation of C69 or G188 had a minimal effect on function, and we observed no effect of mutating U68 or A189. Double mutants that restore the G72·C185, C71·G186 and C70·G187 base-pairing interactions resulted in a complete or nearly complete rescue of the regulatory response, confirming that these base pairs are important for responding to the corrinoid binding state of the aptamer domain (Fig 1F). Together, these results define the functional bases of the kissing loop in this riboswitch as bases C70-C71-G72 in L5 and C185-G186-G187 in L13.

### Dissection of the expression platform of the *P. megaterium metE* riboswitch by mutational analysis

Having established that L13 is part of the kissing loop, we next investigated the mechanism of regulation by alternative RNA conformations in the expression platform by disrupting and restoring predicted base-pairing interactions in the P13, antiterminator, and terminator stems (Figure 2A). We chose to disrupt G-C pairs in the middle of each stem by changing each base to its Watson-Crick-Franklin complement. First, we introduced two C to G point mutations in the 5’ side of the P13 stem. According to the model, mutations at these positions would disrupt the P13 stem, allowing the antiterminator to form and thus preventing stabilization of the terminator. As predicted, these mutations result in constitutive expression (Figure 2A). Next, we introduced two G to C point mutations in the complementary bases on the 3’ side of the P13 stem. In addition to disrupting the P13 stem like the mutations in the 5’ side, these mutations are predicted to disrupt the antiterminator stem, allowing the terminator to form under all conditions. Indeed, this strain has low GFP expression, suggesting the terminator can form even in the absence of corrinoid binding (Figure 2A). We then aimed to restore complementary base-pairing in the P13 stem by combining the mutations in the 5’ and 3’ sides of the P13 stem. As expected, we observed a strong non-inducible phenotype, as this mutant is predicted to be unable to form the antiterminator despite the restoration of base-pairing in the P13 stem (Figure 2A).

**Figure 2.**
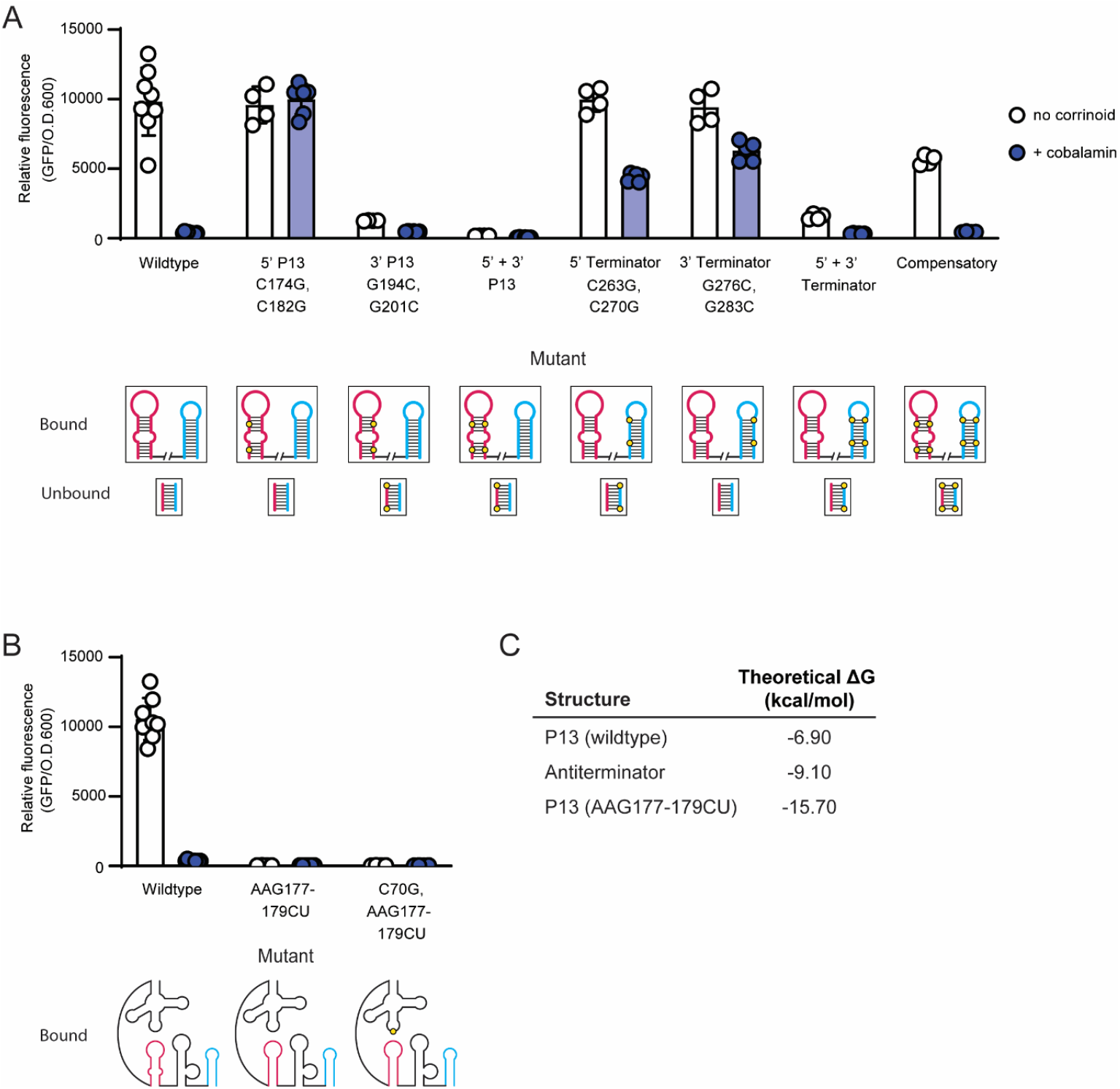
Dissection of the expression platform of the repressing *P. megaterium metE* riboswitch. (A) Influence of point mutations in P13, the terminator, or both stems on gene expression in the *B. subtilis* GFP reporter system without (white) or with (blue) addition of 100 nM cobalamin. The label for each mutant includes the mutated region or the specific mutation, or both. Base numbers are relative to the first base in the P1 stem. For each mutant, a diagram of the P13 (pink) and terminator (blue) hairpins is shown below for the predicted effector-bound conformation and the lower part of the antiterminator stem (pink paired with blue) for the effector-unbound conformation, with the location of each mutation shown as a yellow circle. (B) Phenotypes of mutants designed to close the internal loop in P13. (C) Theoretical ΔG of the wildtype P13, antiterminator, and AAG177-179CU P13 stems calculated in the Structure Editor program (22). Data from four or more biological replicates are shown; bars and error bars represent mean and standard deviation, respectively.

We next made mutations predicted to disrupt the terminator stem. Strains harboring two point mutations in either the 5’ or 3’ sides of the terminator stem were predicted to express GFP constitutively. These strains showed increased expression in the presence of cobalamin, as expected, but retained some inducibility (2.3-fold and 1.5-fold, respectively), suggesting the mutations partially disrupt terminator function (Figure 2A). We then combined the mutations in the 5’ and 3’ sides of the terminator stem, which is predicted to restore complementary base-pairing in the terminator hairpin with the antiterminator stem remaining disrupted, resulting in a non-inducible phenotype. We observed 6-fold reduced expression in the absence of cobalamin, consistent with an inability to form the antiterminator (Figure 2A). These results are consistent with the model shown in Figure 1B.

As an ultimate test of the model for regulation by this riboswitch, we combined the mutations on the 5’ and 3’ sides of P13 and the terminator. This mutant is expected to restore base-pairing in the P13, terminator, and antiterminator stems, and as a result, restore corrinoid-responsive regulatory function. Despite having eight mutations in a structurally complex regulatory domain, this “compensatory” mutant showed an inducible phenotype, indicating restored regulatory function (Figure 2A). This result provides strong evidence in support of our model for the regulatory mechanism of this riboswitch.

### Examining the unpaired regions of the expression platform of the *P. megaterium metE* riboswitch

Our model predicts the presence of an additional structured region containing a large bulge, located between P13 and the terminator in the bound state, which forms the top of the antiterminator in the unbound state (Figure 1B). We found that the large bulge is dispensable for regulation, yet deletion of the entire structured region impacted both repression and maximal expression, suggesting it is important for function (Figure S2).

Our model for corrinoid-responsive regulatory switching relies on the formation of alternative stem-loops in response to the corrinoid-binding state of the aptamer domain (Figure 1B). Implicit in the model is that the antiterminator should be more stable than P13 in the absence of the kissing loop interaction. In this riboswitch, P13 contains an internal loop that we predict sufficiently destabilizes P13 in the absence of the kissing loop to favor formation of the antiterminator stem. To test this aspect of the model, we mutated the 5’ side of the internal loop to bases complementary to those in the 3’ side of the loop, resulting in a closed stem predicted to be more stable than the antiterminator. This mutant showed very low expression (Figure 2B, AAG177-179CU), indicating the stabilized P13 stem prevents antiterminator formation, thus promoting formation of the terminator regardless of the corrinoid-binding state of the aptamer domain. This phenotype is independent of the kissing loop, as disruption of the kissing loop did not influence the phenotype of this mutant (Figure 2B, C70G, AAG177-179CU). Our model for regulatory switching is further supported by calculations of the stability of the wild type and mutant P13 and antiterminator stems. Using the Structure Editor program (22), we estimated the free energy of each predicted stem and found that the antiterminator stem is estimated to be more stable than the wildtype P13 stem, but less stable than the closed AAG177-179CU P13 stem (Figure 2C). Taken together, our mutational analysis of this riboswitch established the interdependent roles of the kissing loop, P13, antiterminator, and terminator stem in regulating gene expression in response to corrinoid binding.

### A model for regulation via a novel activating corrinoid riboswitch

In the course of our study of corrinoid riboswitches from diverse bacteria, we have discovered the first known riboswitch that activates gene expression in response to corrinoids, located upstream of the cobalamin lower ligand activation gene *cobT* in the bacterium *Alkalihalobacillus* (formerly *Bacillus*) *halodurans*. This riboswitch responds to cobinamide (Cbi), a corrinoid lacking a lower ligand, with 8-fold induction in the *B. subtilis* GFP reporter system (Figure 3A). This sequence was previously annotated as a cobalamin riboswitch in a bioinformatic study, but its function has not been experimentally validated (23). We investigated the regulatory mechanism of the *A. halodurans cobT* riboswitch, both to understand how corrinoid binding is coupled to activation and to compare its mechanism with that of the *P. megaterium metE* riboswitch. We propose a model in which L5 and L13 form a kissing loop that stabilizes P13 when a corrinoid is bound to the aptamer domain, as in the *P. megaterium metE* riboswitch (Figure 3B and Figure S1C and D). Unlike the repressing riboswitch, however, P13 and the transcription terminator are mutually exclusive in this model, and thus P13 functions as an antiterminator upon corrinoid binding. We tested the model by introducing mutations predicted to disrupt and restore the kissing loop, P13, and the terminator, using the *B. subtilis* GFP reporter.

**Figure 3.**
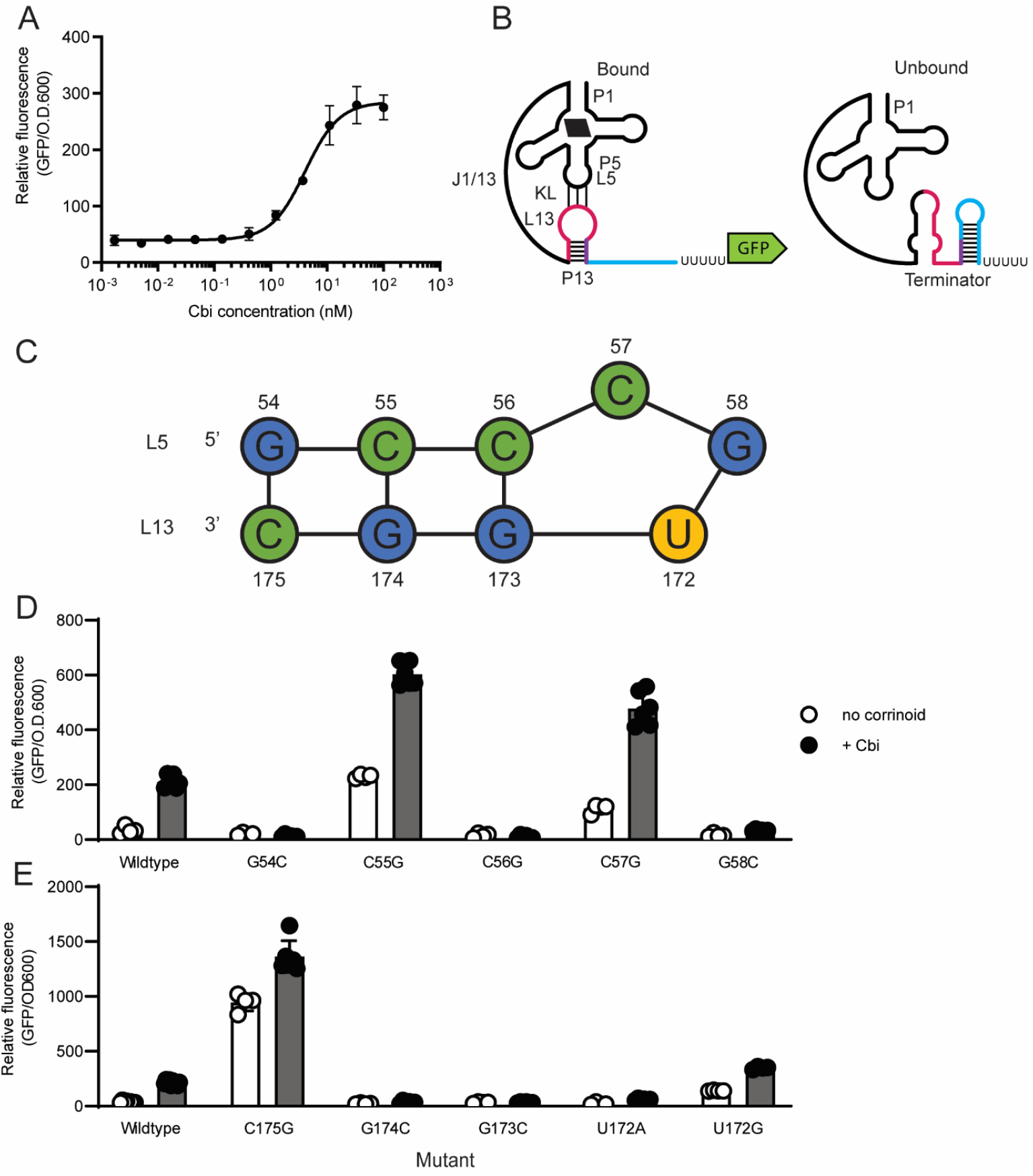
Model for the regulatory mechanism and dissection of the novel activating *A. halodurans cobT* riboswitch. (A) Dose response of the *A. halodurans cobT* riboswitch to cobinamide (Cbi) in the *B. subtilis* GFP reporter. (B) A model for the effector-bound (left) and effector-unbound (right) conformations. The effector-bound state is depicted with Cbi (black parallelogram) and the kissing loop (KL). The color scheme follows that of Fig. 1, but with the region common to P13 and the terminator shown in purple. (C) Diagram of the kissing loop depicting the hypothesized bulge at C57. Base numbers are relative to the first base of P1. The influence of point mutations in (D) L5 and (E) L13 on gene expression was measured in the *B. subtilis* GFP reporter system without (white) or with (blue) addition of 100nM Cbi. Data from four or more biological replicates are shown; bars and error bars represent mean and standard deviation, respectively.

### The activating cobalamin riboswitch relies on a kissing loop

Our model predicts that, like other corrinoid riboswitches, the *A. halodurans cobT* riboswitch relies on a kissing loop for sensing and responding to corrinoids, and therefore disruption of the kissing loop should prevent the riboswitch from activating gene expression. The predicted L5 sequence, GCCCG, is similar to the reported UCCCG consensus sequence. This loop could form base-pairing interactions with four of the ten bases in the predicted L13 sequence, UGGC, with an unpaired C creating a bulge in L5, as observed previously in L13 of the *env8*HyCbl riboswitch (Figure 3C) (17). We found that single point mutations in any of the predicted kissing loop bases disrupted regulatory function, suggesting all are involved in corrinoid-responsive regulation. Mutation of G54, C56, or G58 in L5 or G174, G173, or U172 in L13 to their Watson-Crick-Franklin complement disrupted kissing loop function in the expected way, resulting in non-inducible GFP expression indicative of an inability to sense or to respond to corrinoids (Figure 3D, E). However, mutation of C55 or C57 in L5 or C175 in L13 to their Watson-Crick-Franklin complement, or mutation of U172 to G, resulted in expression levels exceeding that of the wildtype riboswitch, suggesting that the terminator was prevented from forming in these mutants (Figure 3D, E). These are the only mutants with the potential to form four consecutive base pairs, likely a more stable structure than the wildtype kissing loop. Thus, our results suggest that a kissing loop containing four consecutive base pairs stabilizes P13 to the extent that the riboswitch is rarely able to adopt the unbound conformation, similar to the closed-bulge mutant (g) of the *env8*HyCbl riboswitch made by Polaski et al. (17). The 54C-C175G double mutant, which is also predicted to be capable of forming four consecutive base pairs, similarly showed expression levels higher than the wild type (Figure S3). In contrast, double mutants C55G-G174C and C56G-G173C show non-inducible expression despite restoring four base pairs with a bulge, suggesting that both the strength of the kissing loop interaction and the specific bases contained in L5 and L13 contribute to sensing and responding to corrinoid bound by the aptamer domain. Overall, our results support a model in which the bases in L5 and L13 form a kissing loop containing a bulge to stabilize P13 when corrinoid is bound to the aptamer domain and allow the terminator to form when corrinoid is absent (Figure 3B).

### Alternative pairing between bases in P13 and the terminator is responsible for the activating mechanism

We tested this aspect of the model by disrupting and restoring the stems of P13 and the terminator. We found that mutating a single base in the 5’ side of the P13 stem (G168C) results in a non-inducible phenotype, consistent with P13 functioning as an antiterminator (Figure 4). In contrast, changing a single base in the sequence shared by the 3’ side of the P13 stem and the 5’ side of the terminator stem (C181G) results in a constitutive phenotype, as expected, due to disruption of the terminator stem (Figure 4). Disrupting a single base in the 3’ side of the terminator stem (G200C) also results in constitutive expression, but at an expression level 5.3-fold higher compared to disruption of the 5’ side of the terminator, suggesting the sequence context of the terminator influences its strength (Figure 4). The phenotypes of these three single mutants support the hypothesis that P13 and the terminator are alternative secondary structures that inversely influence gene expression.

**Figure 4.**
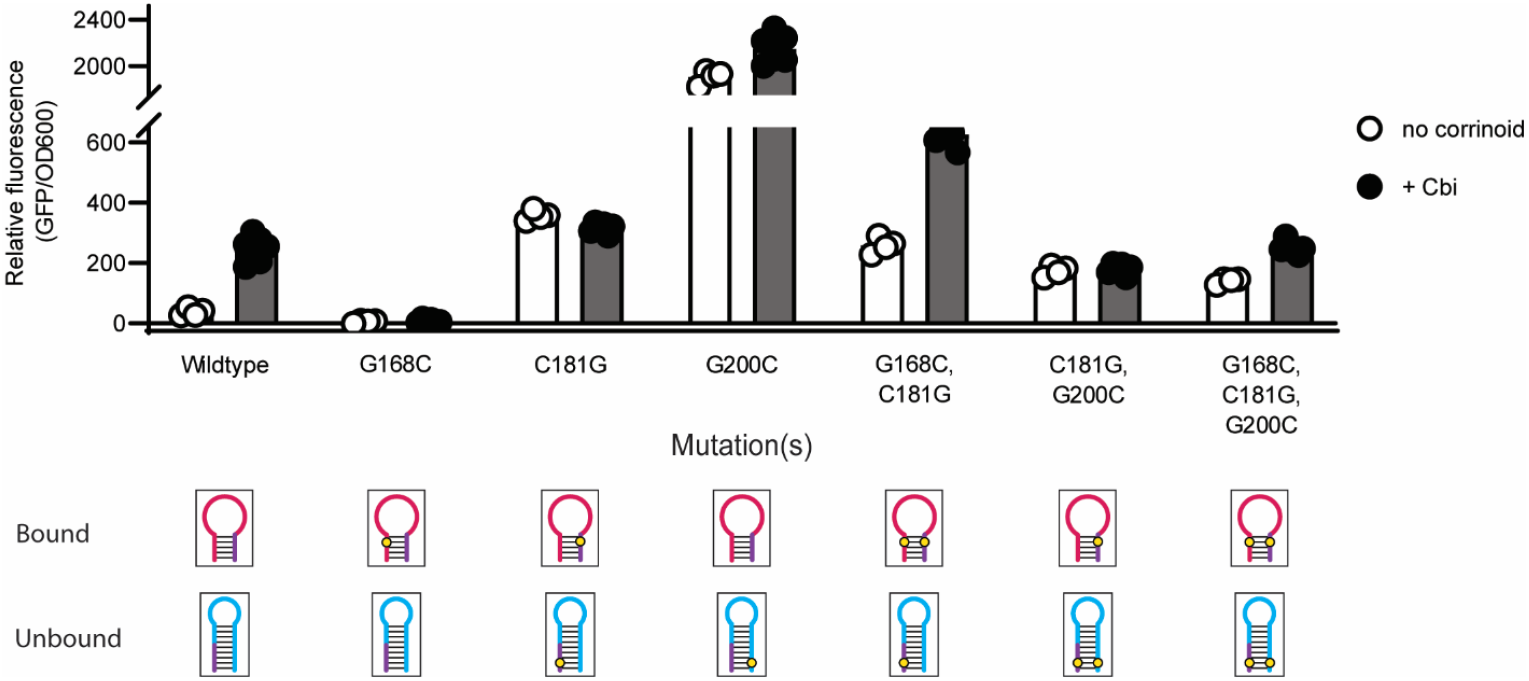
Dissection of the expression platform of the novel activating *A. halodurans cobT* riboswitch. The influence of mutations in P13, the terminator, or both stems on gene expression was measured in the *B. subtilis* GFP reporter system without (white) or with (black) addition of 100nM Cbi. (Bottom) Diagrams of P13 in the predicted effector-bound state and the terminator in the predicted unbound state are shown with the location of each mutation as in Figure 2. The purple region shows the bases that belong to both P13 and the terminator. Data from four or more biological replicates are shown; bars and error bars represent mean and standard deviation, respectively.

The G168C, C181G double mutant was expected to restore the P13 stem but retain the constitutive phenotype of the single C181G mutant due to the disruption of the terminator. This strain shows higher uninduced and induced expression than wild type, with 2.4-fold induction with Cbi addition, suggesting that the terminator retains partial function, allowing some corrinoid-dependent regulation via the restored P13 (Figure 4). The C181G, G200C double mutant was expected to have a non-inducible phenotype due to the restored terminator stem and disrupted P13. This mutant was unable to respond to corrinoid addition, but its intermediate level of expression suggests the restored terminator hairpin is weaker than the wildtype terminator (Figure 4). In the triple G168C, C181G, G200C mutant, nearly 2-fold activation is restored, suggesting that both P13 and the terminator retain partial function in this strain (Figure 4). Overall, these results support the proposed regulatory model, and additionally reveal that both the ability to form alternative structures and the sequences within these structures contribute to the switching function of this riboswitch.

### Corrinoid riboswitches are diverse in sequence and mechanism

Having established and tested models for corrinoid-responsive regulation in one repressing and one activating riboswitch, we sought to understand the extent to which other corrinoid riboswitches may function via the same mechanism. We generated models for the formation of competing structures in the expression platform of two repressing corrinoid riboswitches from *Sporomusa ovata* and tested them by mutational analysis. The *S. ovata cobT* riboswitch responded as predicted when disrupting and restoring P13, but the compensatory mutant did not restore function (Figure S4 and S5). In contrast, the mutations predicted to disrupt and restore regulation in the the *S. ovata nikA* riboswitch did not result in predicted phenotypes, indicating this riboswitch functions via a different mechanism. We hypothesize that multiple alternative base-pairing strategies exist for sensing and responding to corrinoids, due to the remarkable diversity in corrinoid riboswitch sequences (13, 23, 24). This diversity is apparent when comparing the lengths of each subdomain in the 38 corrinoid riboswitches we previously studied in the *B. subtilis* GFP reporter assay (Figure S6) (16). For example, P13 stems range from six to 17 bases in length, and the region between P13 and the terminator, which contains the antiterminator in the *P. megaterium metE* riboswitch, ranges from zero to 82 bases (Figure S6). Thus, it is likely that numerous mechanisms exist for coupling corrinoid binding to gene regulation.

### Flipping the regulatory sign using synthetic expression platforms

A comparison of the regulatory mechanisms for the two riboswitches investigated in this work shows that the main mechanistic difference between the repressing and activating riboswitches is in the nature of the antiterminator: in the repressing riboswitch, it is a structure that forms only when P13 does not form, while in the activating riboswitch the antiterminator is P13 itself. We tested whether these regulatory “rules” can be applied to the design of synthetic riboswitches by attempting to flip the regulatory sign of a repressing or activating riboswitch. In the *B. subtilis yitJ* repressing SAM riboswitch, the regulatory sign was flipped by replacing the expression platform with a modified one from the *B. subtilis pbuE* activating adenine riboswitch (25). However, the kissing loop interaction between the aptamer and expression platform domains in the corrinoid riboswitch makes it less likely that simply exchanging the expression platform will preserve regulatory function.

We constructed a series of engineered expression platforms fused to the aptamer domains of the *P. megaterium metE* or *A. halodurans cobT* riboswitches using two strategies. First, we replaced the entire expression platform of one riboswitch with the other and swapped the sequence in L5 or L13 to preserve the kissing loop interaction. The two chimeric riboswitches designed to activate gene expression in response to corrinoid addition induced GFP expression in response to cobalamin (Figure 5A). Consistent with corrinoid selectivity being encoded in the aptamer domain, these chimeric riboswitches retained selectivity for cobalamin, as they showed little or no response to Cbi (Figure 5A). The two chimeric riboswitches designed to repress GFP expression did not respond to corrinoid addition (Figure S7A).

**Figure 5.**
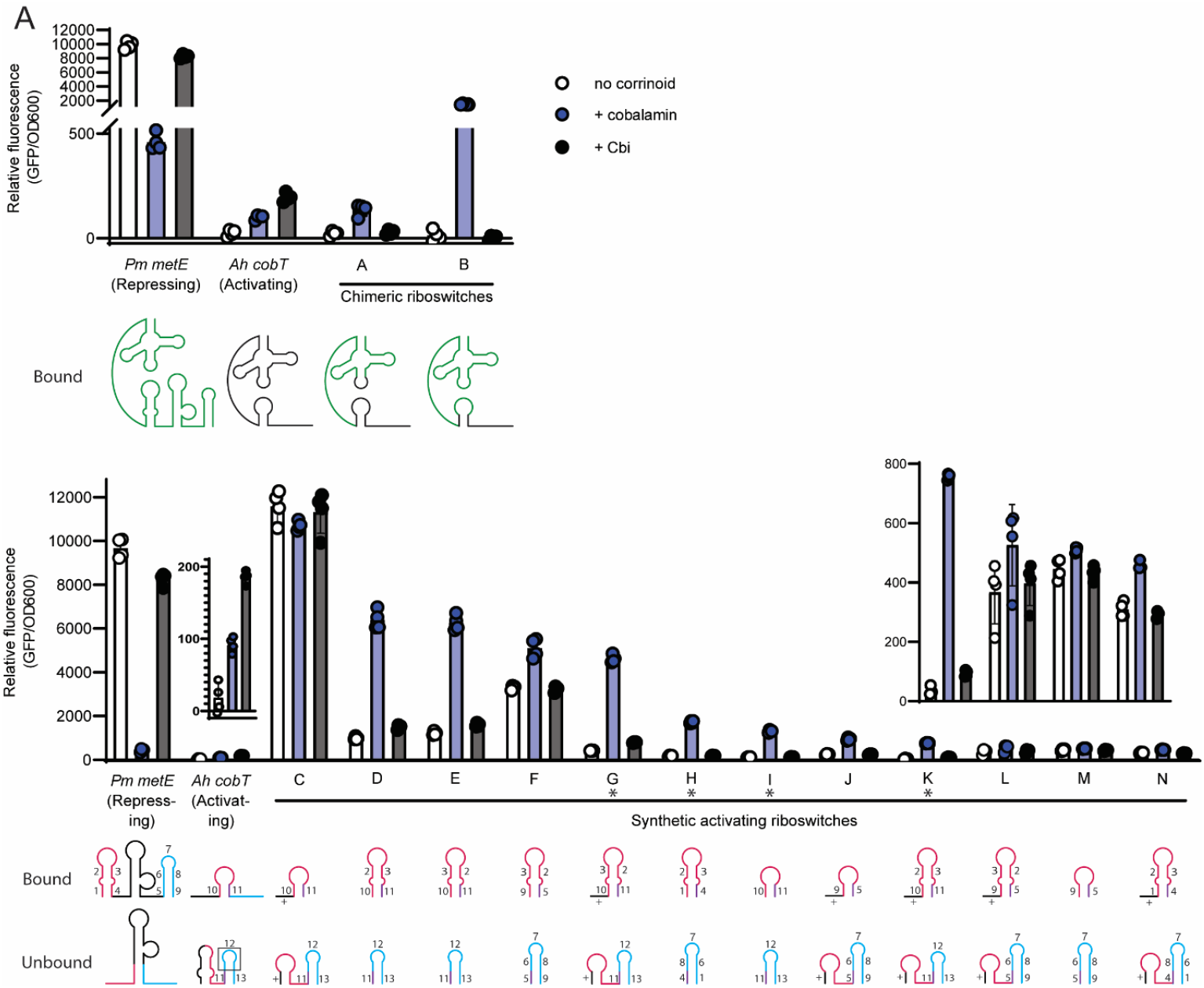
Chimeric and synthetic riboswitches effectively flip the regulatory sign. (A) Chimeric riboswitches were constructed by fusing the *P. megaterium metE* aptamer with the *A. halodurans cobT* expression platform, and gene expression was measured in the *B. subtilis* GFP reporter system with no corrinoid (white), or with addition of 100 nM cobalamin (blue), or Cbi (black). *P. megaterium metE* riboswitch sequences are shown in green and *A. halodurans cobT* sequences in black in the diagrams below, depicting the effector-bound conformation. Kissing loops were preserved by changing either L5 (Riboswitch A) or L13 (Riboswitch B). (B) Synthetic riboswitches were constructed by appending the *P. megaterium metE* aptamer with combined portions of the expression platforms of the *P. megaterium metE* and *A. halodurans cobT* riboswitches. Insets of the wildtype *A. halodurans cobT* riboswitch and synthetic riboswitches K, L, M, and N are shown above the respective strains. Diagrams of the expression platform of each riboswitch construct in the bound (top) and unbound (bottom) conformations are shown below. Numbers represent sequences from P13 (pink and purple) and the terminator (blue and purple) from either the *P. megaterium metE* (numbered 1-9) or *A. halodurans cobT* (10-13) riboswitch. Sequence 12 includes both the loop and part of the terminator stem. Sequences designated with a ‘+’ are entirely synthetic and are the reverse complement of L13 of the *P. megaterium* metE riboswitch. Riboswitches with an asterisk (G, H, I, and K) showed the highest fold change. Data from four biological replicates are shown bars and error bars represent mean and standard deviation, respectively.

In a second strategy, we constructed 20 synthetic expression platforms composed of P13, antiterminator, and terminator hairpins, with different combinations of sequences and lengths. Seven of the 12 synthetic riboswitches designed to activate GFP expression showed induction in response to cobalamin. Notably, these synthetic riboswitches ranged from 8-to over 24-fold induction, higher than in the wildtype *A. halodurans cobT* riboswitch, which was activated 6-fold (Figure 5B). These riboswitches responded only to cobalamin, indicating that, like the chimeric riboswitches, corrinoid selectivity was encoded in the aptamer domain (Figure 5B). There appears to be no correlation between fold induction or expression level and any specific sequence, length of subdomains, or accessory structures among the synthetic riboswitches. Further, none of the synthetic riboswitches designed to convert the *A. halodurans cobT* riboswitch to a repressing riboswitch showed a response to corrinoid (Figure S7B). Nevertheless, these results demonstrate that the mechanistic rules discovered for the activating riboswitch – namely, the formation of alternative structures containing P13 stabilized by the kissing loop in the corrinoid-bound form versus the terminator hairpin in the unbound form – can be applied to design a variety of synthetic riboswitches with higher maximal expression and fold activation than the naturally occurring activating riboswitch of *A. halodurans*.

## Discussion

Riboswitches are remarkable in their ability to undergo conformational changes in response to direct binding to a small molecule. As the first discovered riboswitch, the corrinoid riboswitch has been studied extensively with a range of biochemical techniques, structural approaches, and reporter assays (3, 15–20, 26–28). Here, we gain insight into structural features in the expression platform that influence gene expression by measuring riboswitch function within the natural context of the bacterial cytoplasm. By changing sequences that disrupt or preserve secondary structures, we defined the alternative secondary structures that couple effector binding to transcription in a model repressing riboswitch and a novel activating riboswitch. Applying these themes to the design of synthetic riboswitches showed that new riboswitches can be built using the regulatory scheme we identified.

We present here the discovery and dissection of the first corrinoid riboswitch known to activate gene expression in the presence of a corrinoid. Common to both repressing and activating corrinoid riboswitches are their reliance on a kissing loop to drive formation of alternative secondary structures to sense and respond to corrinoids. Historically, corrinoid riboswitches have been known to repress the expression of genes for corrinoid biosynthesis and uptake to maintain homeostatic intracellular corrinoid levels (4). The presence of an activating corrinoid riboswitch upstream of the corrinoid biosynthesis gene *cobT* in *A. halodurans* diverges from this trend. Riboswitches have been found upstream of *cobT* in many other bacteria, but all of those tested previously repress gene expression upon corrinoid binding (16). *cobT* functions in the late stages of corrinoid biosynthesis by phosphoribosylating the lower ligand base to be attached to Cbi to form a complete corrinoid (29). We hypothesize the difference in regulatory sign between the *A. halodurans cobT* riboswitch and other *cobT* riboswitches lies in their selectivity. Other *cobT* riboswitches tested to date respond most strongly to complete corrinoids (16), which could signal that *cobT* expression is no longer needed and should be repressed. In contrast, the *A. halodurans cobT* riboswitch responds most strongly to Cbi, which is a substrate for enzymes downstream of CobT in the synthesis pathway. Thus, this riboswitch may enable the cell to sense and respond to increased Cbi levels by increasing *cobT* expression in order to complete the final stages of corrinoid biosynthesis.

Overall, our results demonstrate that the main driver of corrinoid riboswitch function is the relative stabilization of alternative secondary structures that promote or prevent transcription elongation. Our results additionally reveal that the sequences within the stems can affect function. For example, changing a single G base on the 5’ side of the terminator of the *A. halodurans cobT* riboswitch affects expression differently from a change in its complement on the 3’ side (Figure 4). We further observed the effect of sequence location when testing synthetic riboswitches G and K (Figure 5B): swapping sequences 2 and 3 in the P13 stem while preserving the same secondary structure and nucleotide content led to differences in expression, again suggesting that the sequence context within hairpins impacts function. The mechanistic basis of these differences should be the subject of future study.

Our experimental results, coupled with the variability in the lengths of hairpins and junctions between hairpins in the expression platform, highlight the versatility of RNA in adopting multiple strategies for achieving the same outcome. The *P. megaterium metE* riboswitch, for example, relies on a large internal structured region between P13 and the terminator for regulatory function. This region of the expression platform is the most variable in length across corrinoid riboswitches, suggesting there are diverse strategies for using alternative secondary structures to regulate expression rather than a single universal mechanism of corrinoid riboswitch regulatory function. Several different models of competing secondary structures in the expression platform have been proposed previously (18–20) and additional mechanisms likely remain to be discovered.

We used the mechanistic rules we uncovered in the mutational analysis of repressing and activating riboswitches to design synthetic riboswitches that convert a repressing riboswitch to an activating riboswitch. The range in corrinoid response in the synthetic riboswitches was surprising, particularly given that they all showed higher maximal expression and most showed higher fold induction than the natural activating riboswitch. Due to their stronger signal, these synthetic riboswitches could potentially be used to detect corrinoids in live cells, food, patient samples, and other samples of interest, or as tools to control gene expression. In light of this, naturally occurring expression platforms can be better utilized as blueprints for engineering precise and robust biosensors and gene regulatory devices.

## Materials and Methods

### Riboswitch sequence manipulation

Secondary structures in the aptamer domain were annotated manually based on the consensus sequence reported in McCown et al. 2017 (13). The P13, terminator, and other stems in the expression platform were annotated using predictions from the StructureEditor program of RNAstructure 6.2 (22). All riboswitch mutant constructs were designed in Benchling. Synthetic expression platforms were designed by combining sequences from the P13 and terminator stems from the *P. megaterium metE* and *A. halodurans cobT* riboswitches. The L13 sequence was sourced from the same riboswitch as the aptamer domain. The P13 stem adopted the five base stem or split ten base stem structure as the two wildtype riboswitches. The terminator was designed to either pair with or overlap with the P13 stem. For the synthetic repressing riboswitches, the 3’ side of P13 paired with the 5’ side of the terminator. For the synthetic activating riboswitches, the 3’ side of P13 shared sequence with the 5’ side of the terminator.

### Strain construction

All *B. subtilis* reporter strains were derived from KK642 (*Em his nprE18 aprE3 eglS*Δ102 *bglT/bglS*ΔEV *lacA*::PxylA-*comK loxP*-Pveg-*btuFCDR queG*::*loxP*) which was derived from strain 1A976 of Zhang et al. (16, 30). All riboswitch mutant constructs were ordered as eBlocks from IDT (Benchling links in Table S1). Each was designed to contain the full-length riboswitch with homology to pKK374 at the NheI (NEB) cut site (16). Linearized pKK374 and the eBlocks were assembled via Gibson assembly. Plasmids were then transformed into XL1-Blue competent cells (UC Berkeley Macrolab) and plated on LB with 100 μg/mL ampicillin. Plasmids from three or four colonies were purified and Sanger sequenced at the Barker DNA Sequencing facility. Plasmids with the correct sequence were linearized with ScaI-HF (NEB) and transformed into the *B. subtilis* fluorescent reporter strain KK642 where they were integrated into the chromosome at the *amyE* locus and plated on LB with 100 μg/mL spectinomycin. Successful integration was confirmed by PCR.

### Riboswitch fluorescent reporter assay

The *B. subtilis* fluorescent reporter strain used in this study and the corrinoid addition assay of riboswitch reporter constructs were developed by Kennedy et al. 2022 (16). Strains containing each riboswitch construct were grown from colonies in LB in a 96-well 2 mL deep well plate and shaken in a benchtop heated plate shaker (Southwest Science) at 37 °C until the cultures reached an optical density at 600 nm OD_600_ of 1.0, usually after 4-5 hours. Cultures were then diluted to a starting OD_600_ of 0.05 into a 96-well microtiter plate (Corning) with either 100 nM cobalamin or Cbi or no corrinoid. Plates were shaken for five hours at 37 °C. A single end point reading of absorbance at 600 nm and GFP fluorescence (excitation/emission/bandwidth = 485/525/10 nm) were measured with a Tecan Infinite M1000 Pro plate reader. Data were plotted and analyzed in GraphPad Prism 9.

## Acknowledgements

We thank Ming Hammond, Karine Gibbs, Kathleen Ryan, Kathleen Collins, Eliotte Garling, Christine Qabar, and all members of the Taga lab for helpful discussions. We thank Zoila Alvarez-Aponte, Zachary Hallberg, Janani Hariharan, Kenny Mok, and Dennis Suazo for critical reading of the manuscript. This work was supported by NIH grant R35GM139633 to M.E.T., NIH grant T32GM132022 to R.R.P. and the Sponsored Projects for Undergraduate Researchers program at UC Berkeley (M.K.). We acknowledge that this work was conducted on the ancestral and unceded land of the Ohlone people.

